# Bacterial community composition of the sea grape *Caulerpa lentillifera*: a comparison between healthy and diseased states

**DOI:** 10.1101/2021.06.30.450479

**Authors:** Germán A. Kopprio, Nguyen Dinh Luyen, Le Huu Cuong, Anna Fricke, Andreas Kunzmann, Le Mai Huong, Astrid Gärdes

## Abstract

The bacterial communities of the sea grape *Caulerpa lentillifera* were studied during a disease outbreak in Vietnam. The Rhodobacteraceae and *Rhodovulum* dominated the composition of healthy *C. lentillifera*. Clear differences between healthy and diseased cases were observed at order, genus and Operational Taxonomic Unit (OTU) level. Bacterial diversity was lower in healthy *C. lentillifera*, probably because of antimicrobial compounds from the macroalgae and/or from *Clostridium, Cutibacterium* or Micrococcus bacteria. The likely beneficial role of *Bradyrhizobium, Paracoccus* and *Brevundimonas* strains on nutrient cycling and phytohormone production was discussed. The white coloration of diseased *C. lentillifera* may not only be associated with pathogens but also with an oxidative response. *Aquibacter, Winogradskyella* and other OTU_s_ of the family Flavobacteriaceae were hypothesized as detrimental bacteria, this family comprises some well-known seaweed pathogens. Moreover, *Thalassobius* OTU 2935 and 1635 may represent detrimental Rhodobacteraceae. *Phycisphaera* together with other Planctomycetes and *Woeseia* were probably saprophytes of *C. lentillifera*. This study offers pioneering insights on the co-occurrence of *C. lentillifera*-attached bacteria, potential detrimental or beneficial microbes, and a baseline for understanding the *C. lentillifera* holobiont. Further metagenomic and biotechnological approaches are needed to confirm functions of some microbes on this macroalgae to enhance food security in the tropics.

## Introduction

The sea grape or green caviar *Caulerpa lentillifera* is a green macroalgae widely cultivated in tropical countries of the Indo-Pacific region, and its market as seafood delicacy has been expanded drastically in the last decades. Not only *C. lentillifera* flavor, vesicular ramuli and succulent texture which resembles caviar, but also elevated contents of polyunsaturated fatty acids, antioxidants, vitamins and trace elements (Saito *et al*., 2010; Paul *et al*., 2014; de Gaillande *et al*., 2017; Yap *et al*., 2019), together with its marketing as functional food increased exponentially its demand. Moreover, the combination of the high nutritional quality of *C. lentillifera* with its simple and low-cost cultivation transforms this macroalgae in a promising candidate to enhance food security of tropical countries (de Gaillande *et al*., 2017; Stuthmann *et al*., 2021). However, one of the major factor contributing to the worldwide decline of macroalgal populations are microbial diseases (Egan *et al*., 2014) and *C. lentillifera* aquaculture is currently endangered by outbreaks.

Macroalgae provide a nutrient-rich niche for microbes and some bacteria are capable to invade algal tissue degrading complex polymers causing disease. Agarases, carrageenases, alginases, fucoidanases, fucanases, mannanases, cellulases and pectinases are depolymerizing enzymes detected in several marine bacteria for the breakdown of algal cell walls (Goecke *et al*., 2010). Moreover, the low genetic diversity of cultured algae and their tight distribution in aquaculture settings facilitate the rapid spread of detrimental microbes. Several bacterial diseases have been reported in commercial macroalgae such as the rote spot and hole-rotten diseases caused by *Pseudomonas* spp. in *Saccharina japonica*, rotten thallus syndrome by *Vibrio* sp. in *Gracilaria verrucosa*, or Anaaki by *Flavobacterium* spp. in *Pyropia yezoensis* (Goecke *et al*., 2010b; Egan *et al*., 2014; Ward *et al*., 2020). Nevertheless, the role of microbes is not only detrimental but can also be beneficial.

Some bacteria are essential for algal health, growth and development. Beneficial bacteria attached to macroalgae induce morphogenesis, fix nitrogen, produce phytohormones, release antifouling and antimicrobial compounds, transport metabolites and nutrients, detoxify pollutants, contribute to algal reproduction and are key for their adaptation to new environments (Aires *et al*., 2013; Hollants *et al*., 2013; Singh and Reddy, 2014). The relation between macroalgae and bacteria is generally mutualistic, and bacteria are mainly benefited with organic substrates for their metabolism and a stable micro-environment protected from predators and changing environmental conditions. The relations between macroalgae and bacterial communities are so tight and reciprocal that the whole entity is considered a holobiont (Egan *et al*., 2013; Arnaud-Haond *et al*., 2017; Califano *et al*., 2020). Furthermore, the bioactive compounds produced by microbes in the holobiont have an enormous potential for drug discovery and biotechnological applications (Friedrich, 2012; Luyen *et al*., 2019).

Despite the commercial importance of *C. lentillifera* and its relevance for human nutrition under future climate-driven and overpopulation scenarios, studies about their microbiome are very limited. Liang *et al*., (2019) described a disease in *C. lentillifera* in Chinese aquaculture systems characterized by a dark-green biofouling, pink-colorated and missing ramuli. In this particular case, Bacteroidetes and Cyanobacteria dominated the bacterial communities of diseased *C. lentillifera*. Although recent advances in the study of the macroalgal microbiome using next generation techniques, the information is still limited and mostly derived from culture methods with their intrinsic limitations. The 16S rRNA gene approach offers broader insights on the natural composition of bacterial communities attached to algae and plays an important role in understanding algal diseases (Egan *et al*., 2014; Kopprio *et al*., 2021). Aims of this work were: 1) to characterize the bacterial communities of *C. lentillifera* in Vietnamese aquaculture, and 2) to assess the effect of an unknown disease on its bacterial community composition. We hypothesize strong differences in bacterial communities of *C. lentillifera* between healthy and diseased cases.

## Results

The order Rhodobacterales (class Alphaproteobacteria) and the genus *Rhodovulum* of the mentioned order presented the highest relative sequence abundance in *C. lentillifera* at each respective taxonomic level (Fig. 1). The Rhodobacteraceae was the only family detected within the order Rhodobacterales. According to the Mann-Whitney test, the mean relative abundance of Clostridiales (phylum Firmicutes) was significantly higher in healthy *C. lentillifera*, whereas the abundances of Flavobacteriales (phylum Bacteroidetes), Phycisphaerales (phylum Planctomycetes) and Steroidobacterales (class Gammaproteobacteria) characterized diseased *C. lentillifera* thalli. The 99.7 % of the OTUs for the order Flavobacteriales corresponded to the family Flavobacteriaceae. The trend observed at order level was similar at genus level: the relative sequence abundance of *Clostridium sensu stricto* 7 (Clostridiales) was significantly higher in healthy cases, while *Aquibacter* and Flavobacteriaceae unclassified (Flavobacteriales), *Phycisphaera* (Phycisphaerales) and *Woeseia* (Steroidobacterales) were typical of diseased cases. Without significant differences, other abundant genera were *Rhodovulum, Cutibacterium* and *Paracocccus* for healthy *C. lentillifera*, while *Thalassobius, Tropicibacter* and *Blastopirellula* for diseased *C. lentillifera*.

**Fig 1.**
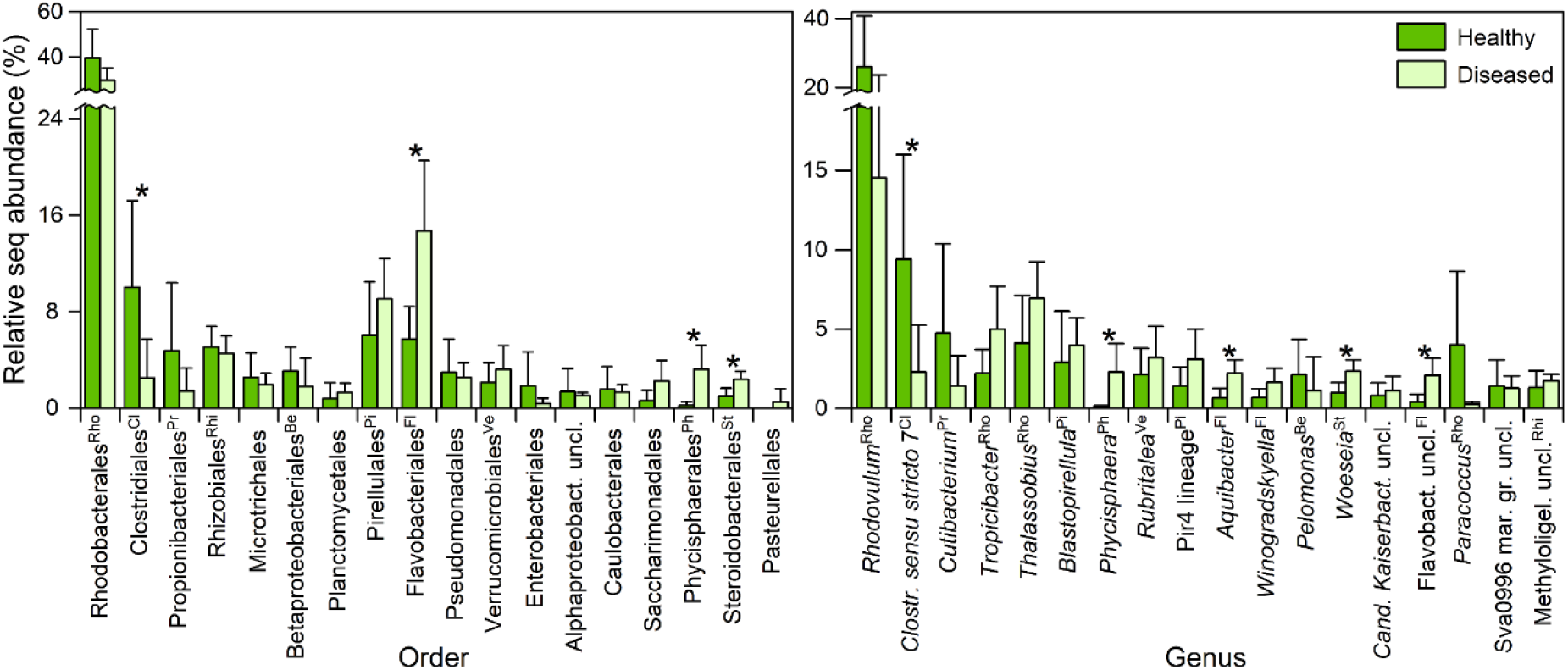
Mean relative sequence abundance (± standard deviation) of the main orders and genera in healthy and diseased individuals of the sea grape *Caulerpa lentillifera*. (^*^) indicates significant differences at p < 0.05 according to Mann-Whitney test. Caption superscript: shows correspondence between order and genus (e.g., ^Rho^ = Rhodobacterales). uncl.: unclassified, Aphaproteobact.: Alphaproteobacteria, *Clostr*.: *Clostridium, Cand. Kaiserbact*.: *Candidatus Kaiserbacteria*, Flavobact.: Flavobacteriaceae, mar. gr.: marine group, Methyloligel.: Methyloligellaceae

PERMANOVA revealed significant differences between the healthy and diseased cases at order level (Pseudo-F = 3.89, p = 0.016). According to SIMPER analyses, the total dissimilarity between healthy and diseased cases was 41.3%. The orders which presented significant differences in the Mann-Whitney test were also important contributors to the dissimilarities and evidenced the same trend in healthy or diseased *C. lentillifera* (Table I). Other relevant orders with higher percentages of dissimilarities but without significant differences were Propionibacteriales and Betaproteobacteriales for the healthy cases, while Saccharimonadales and Verrucomicrobiales for the diseased cases. Furthermore, PERMANOVA detected significant differences at OTU level between healthy and diseased *C. lentillifera* (Pseudo-F = 3.12, p = 0.015). According to SIMPER analysis, the total dissimilarity at OTU level was 71.3% and the main OTUs contributing to this value are detailed in Table I.

**Table I.**
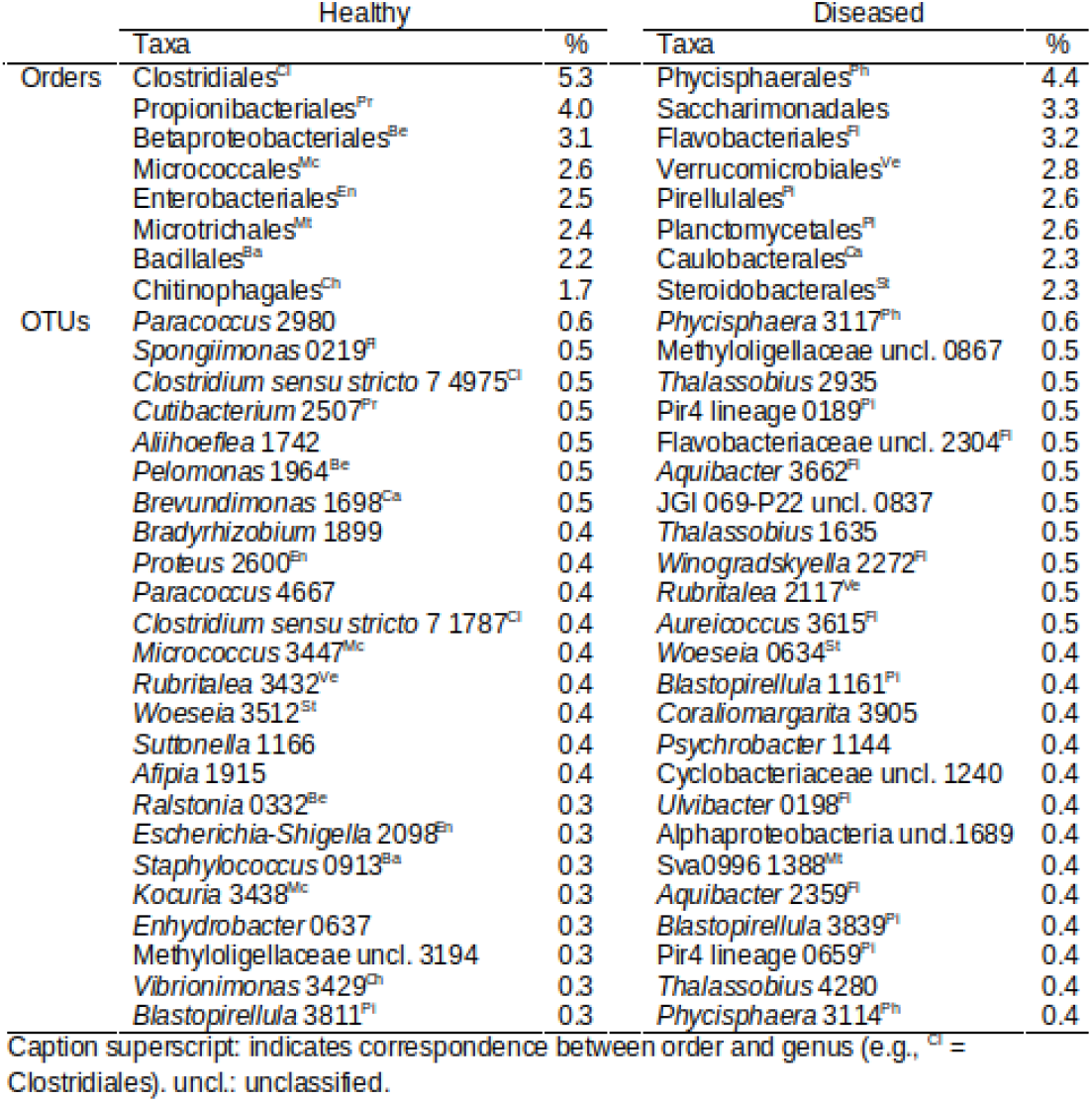
Main orders and operational taxonomic units (OTUs) of the sea grape *Caulerpa lentillifera* contributing to the dissimilarity (41 and 71 % in total, respectively) according to SIMPER analyses. Taxa are grouped based on their dominance in healthy or diseased cases.

The OTUs of order Clostridiales, typical of healthy *C. lentillifera*, were *Clostridium sensu stricto* 7 4975 and 1787 (Table I). Other OTUs dominating with elevated percentages of dissimilarity in the healthy cases were those of the genera: *Paracoccus, Spongiimonas, Cutibacterium, Aliihoeflea, Pelomonas, Brevundimonas* and *Bradyrhizobium*. In the case of diseased cases, the major OTUs were those of Flavobacteriaceae unidentified, *Aquibacter, Winogradskyella, Aureicoccus* and *Ulvibacter* for the order Flavobacteriales (all belonging to the family Flavobacteriaceae), *Phycisphaera* for the order Phycisphaerales and *Woeseia* for the order Steroidobacterales. Other OTUs with elevated percentages of dissimilarity in the diseased cases were Methyloligellaceae unclassified 0867, *Thalassobius* 2935, Pir4 lineage 0189 and JGI 069-P22 unclassified 0837. A list with other relevant OTUs according to their dissimilarity is presented in table I. Moreover, the NMDS ordination of *C. lentillifera* samples at OTU level (Fig 2) suggested contrasting bacterial communities between healthy and diseased cases.

**Fig 2.**
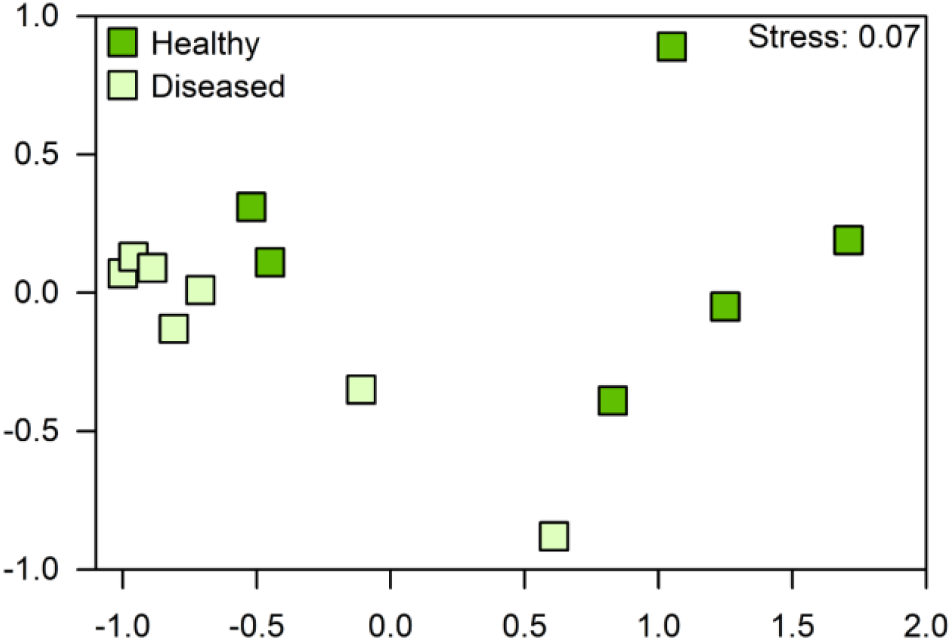
Non-metric Multi-Dimensional Scaling (NMDS) ordination of healthy and diseased individuals of the sea grape Caulerpa lentillifera at Operational Taxonomic Unit (OTU) level

The heat map (Fig. 3) showed a similar cluster of 4 diseased *C. lentillifera*, while a variable ordination of the other cases. OTUs associated exclusively to diseased status were *Aquibacter* 3662, Flavobacteriaceae unclassified 2304, *Woeseia* 0634, *Phycisphaera* 3117 and *Muricauda* 2354; while those exclusively detected in the healthy status were *Bradyrhizobium* 1899, *Paracoccus* 4667 and *Paracoccus* 2980. OTUs of the genera JGI 069-P22 unclassified, *Thalassobius*, Saccharimonadales unclassified, Flavobacteriaceae unclassified, *Winogradskyella, Aureicoccus*, Pir4 lineage, *Tropicibacter, Algisphaera*, Cellvibrionaceae unclassified and *Aquibacter* dominated mainly in the diseased cases. Other OTUs which dominated but not exclusively in the healthy cases are displayed in the heat map. The inverse Simpson indexes at the bottom of the heat map indicated mainly a higher diversity in diseased *C. lentillifera*.

**Fig 3.**
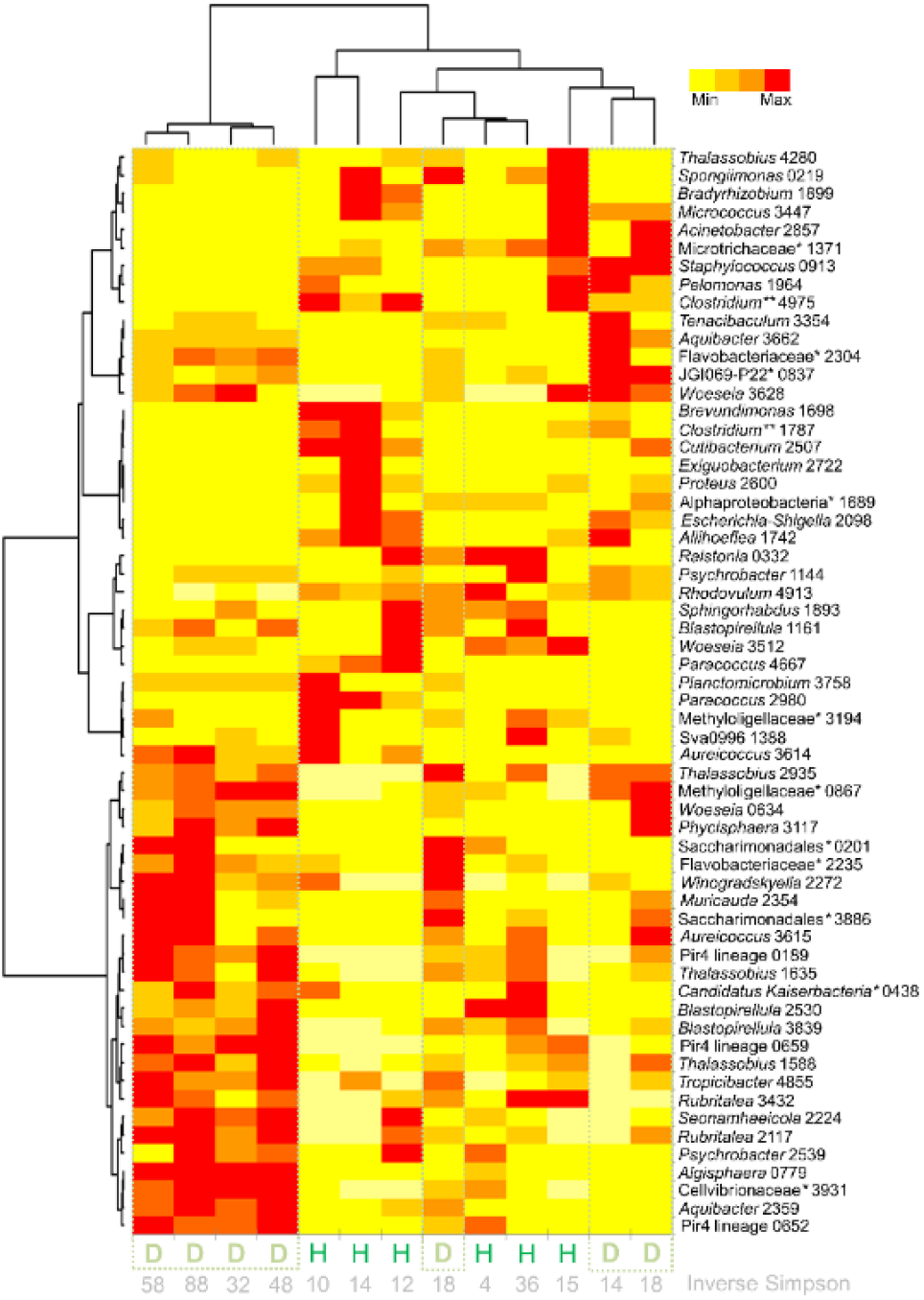
Heatmap of the main Operational Taxonomic Units (OTUs) on healthy (H) and diseased (D) individuals of the sea grape *Caulerpa lentillifera*. (^*^) unclassified (^**^) *sensu stricto* 7

## Discussion

### Main composition and potential beneficial bacteria

The bacterial communities of *C. lentillifera* presented a similar composition than other green algae at higher taxonomic levels. The dominance of Alphaproteobacteria and Bacteroidetes has been reported in green macroalgae and some *Caulerpa* species (Friedrich, 2012; Aires *et al*., 2013; Singh and Reddy, 2014). The order Rhodobacterales and family Rhodobacteraceae dominated the bacterial communities of *C. lentillifera* in this particular case from the Vietnamese aquaculture. Bacteria within the order Rhodobacterales are generally surface colonizers and facilitate the settlement of microbial communities with the production of extracellular polymeric substances (Kopprio *et al*., 2020). According to Simon *et al*. (2017), marine Rhodobacteraceae are characterized by genes for the degradation of sulfated polysaccharides from algae, production of phytohormones, metabolism of osmolytes, transport of metals and detoxification.

The bacterial communities at lower taxonomic levels are generally highly variable in seaweeds and their composition may be associated with a particular function within the holobiont. The microbiome composition depends generally on the geographic location, environmental factors, seaweed tissues, developmental stage and even among local individuals (reviewed by Friedrich, 2012). Burke *et al*., (2011) explain the pattern of bacterial colonization on a green macroalgae with the competitive lottery model, according to this model different species with similar functional traits can occupy the same niche depending on a stochastic probability. The highest relative abundance of Rhodobacterales coincided with the results obtained in *C. lentillifera* from China (Liang *et al*., 2019); nevertheless, one of the differences was in the dominant genus within this order: *Leisingera* in contrast to *Rhodovulum* in our study. Moreover, the last authors reported a higher abundance Planctomycetales and Oceanospirillales in healthy *C. lentillifera* and not the elevated values of Clostridiales observed in this study.

*Rhodovulum* and particularly *Paracoccus* are Rhodobacteraceae with potential beneficial roles for *C. lentillifera. Rhodovulum* species have a high metabolic versatility with the capability to degrade organic pollutants, to produce polymers and to contribute with photosynthetic functions (Khandavalli *et al*., 2018; Foong *et al*., 2019; Baker *et al*., 2021). *Paracoccus* is a common genus among seaweed-associated bacteria with a key role in the cycle of nitrogen and in the production of siderophores (Mei *et al*., 2019). *Paracoccus* strains promote the growth and the morphogenesis in *Ulva* species (Ghaderiardakani *et al*., 2017) and were described as auxin producers (Kurepin *et al*., 2014). Plant hormone production seems to be widespread in various genera of marine bacteria (Goecke *et al*., 2010). Furthermore, *Paracoccus* sp. presented algicidal activity (Zhang *et al*., 2018) and some members of this genus might contribute with antifouling activities. The macroalgal surfaces constitute a highly competitive niche for nutrient and space.

The likely production of antimicrobial compound of *C. lentillifera* may explain partially the low diversity in healthy *C. lentillifera*. The same diversity trend was reported comparing healthy and diseased *C. lentillifera* from the Chinese aquaculture (Liang *et al*., 2019). Some *Caulerpa* species contain antibacterial compounds such as alkaloids, terpenoids, phenols and flavonoids (Goecke *et al*., 2010; Yap *et al*., 2019; Zainuddin *et al*., 2019). Moreover, strains of *Clostridium sensu stricto* 7 and *Cutibacterium* may reduce diversity with the production of antimicrobial metabolites in healthy *C. lentillifera*. Some green algae provide a habitat for *Clostridium* species and even for potential pathogens (Chun *et al*., 2013). *Clostridium* spp. can catabolize several algal polymers (Song *et al*., 2011), have a likely function of copper detoxification in *Codium tormentosum* (Le Pennec and Gall, 2019), and are sources of novel antimicrobial compounds (Schieferdecker *et al*., 2019; Pahalagedara *et al*., 2020). A new thiopeptide antibiotic as microflora modulator was detected on *Cutibacterium* sp. (Claesen *et al*., 2020) and some species of this genus might produce antibiotics on *Kappaphycus striatus* (Kopprio *et al*., 2021). A similar role on algal defense is suspected in the other Actinobacteria *Micrococcus* spp. (e.g., Hollants *et al*., 2013). In addition, this genus together with *Bradyrhizobium* were reported as beneficial endophytic bacteria in many terrestrial plants with agronomic value (Afzal *et al*., 2019).

*Bradyrhizobium* and *Allihoeflea* of the order Rhizobiales may participate in the cycle of nitrogen and the production of phytohormones. “Rhizobacteria” were extensively studied in terrestrial plants because of their ability to fix nitrogen and to modify genetically their host. Some organisms of the family Rhizobiaceae presented growth-enhancing and probiotic properties on green microalgae (Rivas *et al*., 2010). Furthermore, an endosymbiotic bacterium of this taxa was responsible for the nitrogen supply in *Caulerpa taxifolia* (Chisholm *et al*., 1996).

*Bradyrhizobium japonicum* increases the biomass and starch content of the green microalgae *Chlamydomonas reinhardtii* (Xu *et al*., 2016) and some strains of *B. japonicum* are producers of the several phytohormones such as indole-3-acetic acid, gibberellic acid, zeatin, abscisic acid and ethylene (Boiero *et al*., 2007).

A key role for *C. lentillifera* growth and health is supposed on strains of *Brevundimonas* spp. Member of this genus establish a symbiotic relationship with green microalgae promoting their growth, producing indole-3-acetic acid and enhancing nutrient uptake (Tate *et al*., 2013; Sforza *et al*., 2018; Zhang *et al*., 2021). Some of their strains have a potential antifouling activity against cyanobacteria (Lin *et al*., 2014) and in some terrestrial plants: alleviate toxicity, fix nitrogen and promote growth (Singh *et al*., 2016; Naqqash *et al*., 2020). *Pelomonas* OTU 1964 sustains presumably a healthy status on *C. lentillifera*. This genus is common in the rhizosphere of some plants with a nitrogen fixation function (Terakado-Tonooka *et al*., 2008) and some of their aquatic strains produce the antibacterial compounds such as pelopuradazole (He *et al*., 2014). Surprisingly, potential human pathogens like *Escherichia-Shigella* and *Staphylococcus* were related to the healthy state of *C. lentillifera* according to SIMPER analysis. Hollants *et al*., (2013) reported these bacteria with the beneficial functions of defense and morphogenesis in seaweeds, respectively.

### Potential detrimental bacteria

The lack of production of antibacterial compounds by diseased *C. lentillifera* or the absence of of some defense microorganisms may increase bacterial diversity. As mentioned in the above section, this pattern was also observed in China, where the orders Flavobacteriales, Phycisphaerales and Cellvibrionales dominated the diseased and decayed thalli of *C. lentillifera* (Liang *et al*., 2019). With different disease symptoms, a similar pattern was observed in our study: Flavobacteriales, Phycisphaerales and an unidentified genus of the family Cellvibrionaceae were typical of diseased *C. lentillifera*. The bleaching and white coloration symptom in *C. lentillifera* may not only be linked to a pathogen invasion, but also to an oxidative burst response as a defense mechanism. Some *Caulerpa species* produce reactive oxygen species like hydrogen peroxide against invasive organisms (Box *et al*., 2008).

The copiotrophic nature and ability to catabolize algal polymers of the Flavobacteriaceae gives to this family, the potential to breakdown algal walls, to invade tissues and to cause disease under certain conditions. Bacteroidetes and particularly Flavobacteriaceae are degraders of complex biopolymers like sulfated polysaccharides from algal tissues (Yilmaz *et al*., 2016; Jain *et al*., 2019). Many members of the Flavobacteriaceae have been confirmed or suspected as causative agents of seaweed diseases (Goecke *et al*., 2010; Kumar *et al*., 2016; Ward *et al*., 2020). In our particular case, we hypothesize as potential pathogens: strains of *Aquibacter, Winogradskyella, Aureicoccus, Ulvibacter* and *Muricauda. Aquibacter* spp. characterized the senescent state and decayed phase of the diatom *Skeletonema dohrnii* (Liu *et al*., 2021). In the case of *Winogradskyella spp*., higher abundances have been observed on *Pyropia yezoensis* with yellow spot disease (Liu *et al*., 2020) and on *Delisea pulchra* with a bleaching disease (Kumar *et al*., 2016). On the other hand, Flavobacteriales are a common order among green algae (Hollants *et al*., 2013) and *Spongiimonas* was linked to healthy *C. lentillifera*.

We hypothesize that *Phycisphaera* and *Algisphaera* spp. of the order Phycisphaerales were opportunistic pathogens or saprophytes of *C. lentillifera*. As mentioned before, the order Phycisphaerales was typical of the disease cases from the Chinese aquaculture and strains of these genera were isolated from marine macroalgae presenting agarolitic activity (Fukunaga *et al*., 2009; Yoon *et al*., 2014). Nevertheless, Planctomycetes are widespread in the epiphytic microbial community of macroalgae, possess potential beneficial effects for their host like the production of bioactive compounds, and are rarely identified as seaweed pathogens (Lage and Bondoso, 2014). Some strains of *Phycisphaera, Algisphaera*, Pir 4 lineage and *Blastopirellula* may be using their higher number of sulfatases for the degradation of algal polysaccharides (Bondoso *et al*., 2017) from the dead or senescent tissue of *C. lentillifera*, which could be injured by other pathogen or degraded because of an oxidative defense response. Other probable saprophyte was *Woeseia*, species of this genus can utilize a broad range of energy-yielding metabolisms and substrates (Mußmann et al., 2017). Some bacteria may be specialized on small molecules derived from the breakdown of algal polymers and benefited from the degradation of algal tissue. Members of the family Methyloligellaceae oxidize generally one carbon atom molecules (Walker *et al*., 2021) and Patescibacteria (JGI 069-P22 and some Saccharimonadales in this study) may be using monosaccharides because they have reduced genes for polysaccharide catabolism in their simple genomes (Tian *et al*., 2020).

The classification of potential beneficial or detrimental bacteria at higher taxonomic levels in *C lentillifera* is not clear. The orders Verrucomicrobiales is well-known as active polysaccharide degraders (Martinez-Garcia *et al*., 2012) but only a few genera were linked to algal disease. For example, several OTUs of *Rubritalea* within this order were typical of *Saccharina japonica* with symptoms of rotten hole disease (Zhang *et al*., 2020). Furthermore, the ability to metabolize algal compounds of Rhodobacteraceae in a mutualistic relationship with seaweeds, may became detrimental with some members of this family such as *Thalassobius* and *Tropicibacter. Thalassobius* spp. characterized diseased *Delisea pulchra* (Kumar *et al*., 2016) and were responsible of the direct lysis of a red-tide microalgae (Wang *et al*., 2010). *Tropicibacter multivorans* has been detected in the microbiome of *Caulerpa cylindraceae* but little information is available about its function (Rizzo *et al*., 2016). In this study case, *Tropicibacter* spp. may contribute with the degradation of algal polymers.

## Conclusions

Despite of different symptoms comparing cases from Vietnamese and Chinese *C. lentillifera* cultures, a common bacterial pattern was observed on diseased organisms. A disease on *C. lentillifera* may be caused by a community of detrimental bacteria together with changes on algal response, and not only by a particular pathogen. Moreover, with this 16S rRNA approach is not possible to identify clearly pathogens from saprophytes. A mutualistic relationship with a particular bacteria may change and became detrimental under certain conditions; nevertheless, clear differences between the healthy and diseased states were observed at several taxonomic levels. This work explores changes in the bacterial community composition of *C. lentillifera* under two conditions, detects co-occurrence but not causality, we cannot discard other etiological agent not covered by the selected amplicon. Common patterns and roles were inferred on this pioneer work from the Vietnamese aquaculture, which contribute to the understanding of potential key players in the holobiont. A further metagenomic approach will confirm the presence of functional genes for a particular microorganisms and their potential biotechnological applications.

## Experimental Procedures

### Culture conditions, sampling and disease symptoms

*Caulerpa lentillifera* thalli were sampled in the culture facilities of the company VIJA at Van Phong bay (N 12° 35’ 17.67”, E 109°13’ 39.76”) in the central eastern coastline of Vietnam. The ponds for *C. lentillifera* culture were used previously for shrimp farming and contained apparently a considerable load of organic matter. Water temperature in the ponds reached ∼31°C in May, with pH values of ∼ 8.1, conductivity ∼53 mS cm^-1^ and dissolved oxygen ∼5.6 mg L^-1^. The sea grape ponds were covered from direct sunlight radiance with shade cloths. Six healthy and seven diseased thalli of *C. lentillifera* were collected during an outbreak of an unknown disease. In contrast to the bright green color of healthy individuals (Fig. 1A), the diseased *C. lentillifera* showed a marked tissue discoloration and a white biofilm on the external circumference of the vesicular ramuli (Fig. 1B). After the described symptoms, the diseased algae lost their turgid appearance and decayed with the consequent loss of their commercial value.

**Fig 4.**
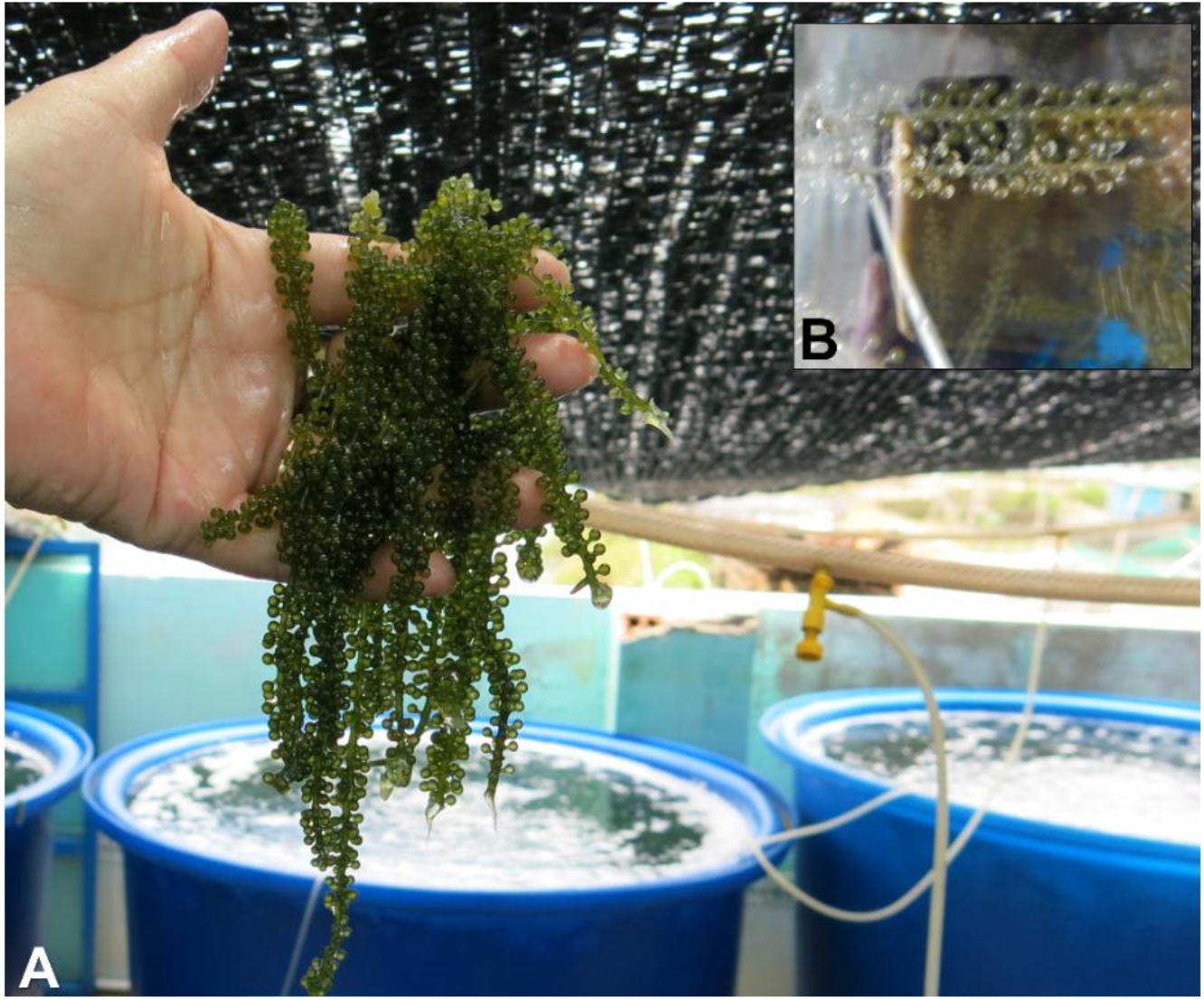
A) Healthy thalli of *Caulerpa lentillifera* and aquaculture facilities in Vietnam. B) A comparison between healthy (up) and diseased (down) individuals, the diseased individuals showed bleaching of thalli and a white biofilm on the external circumference of the ramuli.

### DNA extraction and amplification

*Caulerpa lentillifera* individuals were collected in sterile vials and preserved at -20°C until further analysis. Under laboratory conditions, DNA was extracted basically according to Griffiths *et al*., (2000). Fragments of the thalli were immersed in 600 µL of hexadecyltrimethylammonium bromide (CTAB) with 60 µL of 10 % SDS and 60 µL of 10 % N-lauroylsarcosin. Glass beads were added and the tissue was homogenized in a FastPrep at 5 m s^-1^. Homogenates were rinsed with 600 µL of phenol-chloroform-isoamylalcohol (25:24:1) and centrifuged at 16,000 g and 4°C for 10 min. The upper aqueous layer was transferred to a clean reaction tube, mixed with 2 volumes of 30 % polyethylene glycol (PEG) 6000 and 1.6 M NaCl and incubated at 4°C for 120 min. Subsequently, the sample was centrifuged at 17,000 g and 4°C for 90 min, the supernatant was carefully removed and the pellet was washed with ice-cold 70% ethanol. The pellet was air-dry at 37°C for a few minutes, dissolved in PCR grade water and stored at -20°C. The hypervariable region V3-V4 of the 16S rRNA gene was amplified with the set of primers according to Klindworth *et al*., (2013): Bact-341F (5′-3′: CCT ACG GGN GGC WGC AG) and Bact-785R (GAC TAC HVG GGT ATC TAA KCC)

### Sequencing, bioinformatics and statistical analysis

The amplicon V3-V4 of the 16S rRNA gene was sequenced with a 2 × 300-bp paired-end run on an Illumina MiSeq platform. Sequencing, removal of primer sequences and demultiplexing were conducted by the company LGC genomics. Sequences were trimmed and merged using Trimmomatic v0.36 (Bolger *et al*., 2014) and PEAR v0.9.8 (Zhang *et al*., 2014), respectively. Operational Taxonomic Units (OTUs) were clustered by Minimum Entropy Decomposition MED v2.1 (Eren *et al*., 2015) and their representatives were submitted to SilvaNGS (v132, https://ngs.arb-silva.de/silvangs/) using a sequence similarity of one for clustering and the remaining variables as default. Singleton, doubleton and sequences from mitochondria, chloroplasts and archaea were removed from the analysis. A total of ∼222,000 reads remained after data curation and the mean OTU number per sample was ∼17,100 ± 14,500 (± standard deviation). Demultiplexed and primer-clipped sequences were deposited at the European Nucleotide Archive (ENA) using the data brokerage service of the German Federation for Biological Data (Diepenbroek *et al*., 2014) with the number PRJEB42826, in compliance with the Minimal Information about any (x) Sequence (MIxS) standard (Yilmaz *et al*., 2011).

Pooling of taxa, removal of OTUs with poor alignment quality and relative sequence abundance and diversity calculations were conducted in R v4.0.5 (R Core Team, 2021) and additional packages. The relative sequence abundances of the main orders between the diseased and healthy cases were compared with a Mann-Whitney test. Data were transformed, a Bray Curtis similarity matrix was calculated, and differences between the states at order and OTU level were evaluated using permutational multivariate analysis of variance (PERMANOVA). In case of significance differences, similarity percentage analysis (SIMPER) was performed to detect the main taxa contributing to dissimilarities. Samples at OTU level were ordinated by Non-metric Multi-Dimensional Scaling (NMDS). A heat map based on the 60 most abundant OTUs, covering the 64 ± 11 % of the total relative sequence abundance, was displayed to compare healthy and diseased cases. Graphics and statistics were performed with R 4.0.5, Xact 7.21, PRIMER v6 + PERMANOVA, and XLSTAT-Ecology

## Acknowledgment

We are grateful with T. M. Duc for logistical support.

## Data availability

16S sequencing data are available at ENA (https://www.ebi.ac.uk/ena/data/view/PRJEB42826).

